# Beach sand beneath our feet: an overlooked reservoir of antibiotic resistance genes and pathogens, revealed by metagenomic evidence from Qingdao’s recreational beaches

**DOI:** 10.1101/2025.06.25.661489

**Authors:** Chenglu Li, Hongyue Ma, Yidan Chang, Hezi Ge, Xiaoyun Liu, Jiangtao Xie, He Zhang, Pengfei Cui

## Abstract

Coastal beach environments, especially those with heavy tourist activity, represent a dynamic interface between terrestrial and marine microbiomes. However, beach sand is rarely considered in surveillance of antibiotic resistance. In this study, we combined 16S rRNA gene sequencing and shotgun metagenomics to characterize microbial communities and antibiotic resistance genes (ARGs) at four popular recreational beaches in Qingdao, China. We show that beach sands harbor highly diverse bacterial communities with significantly greater richness than adjacent seawater. Beach sand near sewage discharge points had microbial profiles intermediate between raw sewage and open-ocean water, reflecting contributions from both sources. Metagenomic analysis identified a broad “resistome” of over 300 distinct ARGs in these beach and water samples, spanning 33 antibiotic classes and 6 resistance mechanism categories. Notably, the rifampicin resistance gene *rpoB2* was the most abundant ARG, and genes conferring resistance to peptide antibiotics were prevalent. Sewage outfalls were found to be major contributors of ARGs, with waste-influenced sand containing a larger and more diverse ARG pool than sand farther from pollution sources. We also detected a high diversity of bacterial virulence factor genes, particularly in sand. Our findings highlight that tourist beaches can serve as reservoirs and mixing zones for antibiotic-resistant and potentially pathogenic bacteria from land-based pollution. This underscores the need to include beach sand in environmental monitoring for antibiotic resistance and to improve wastewater management at recreational coastlines to mitigate public health risks.

**Graphical abstract:** 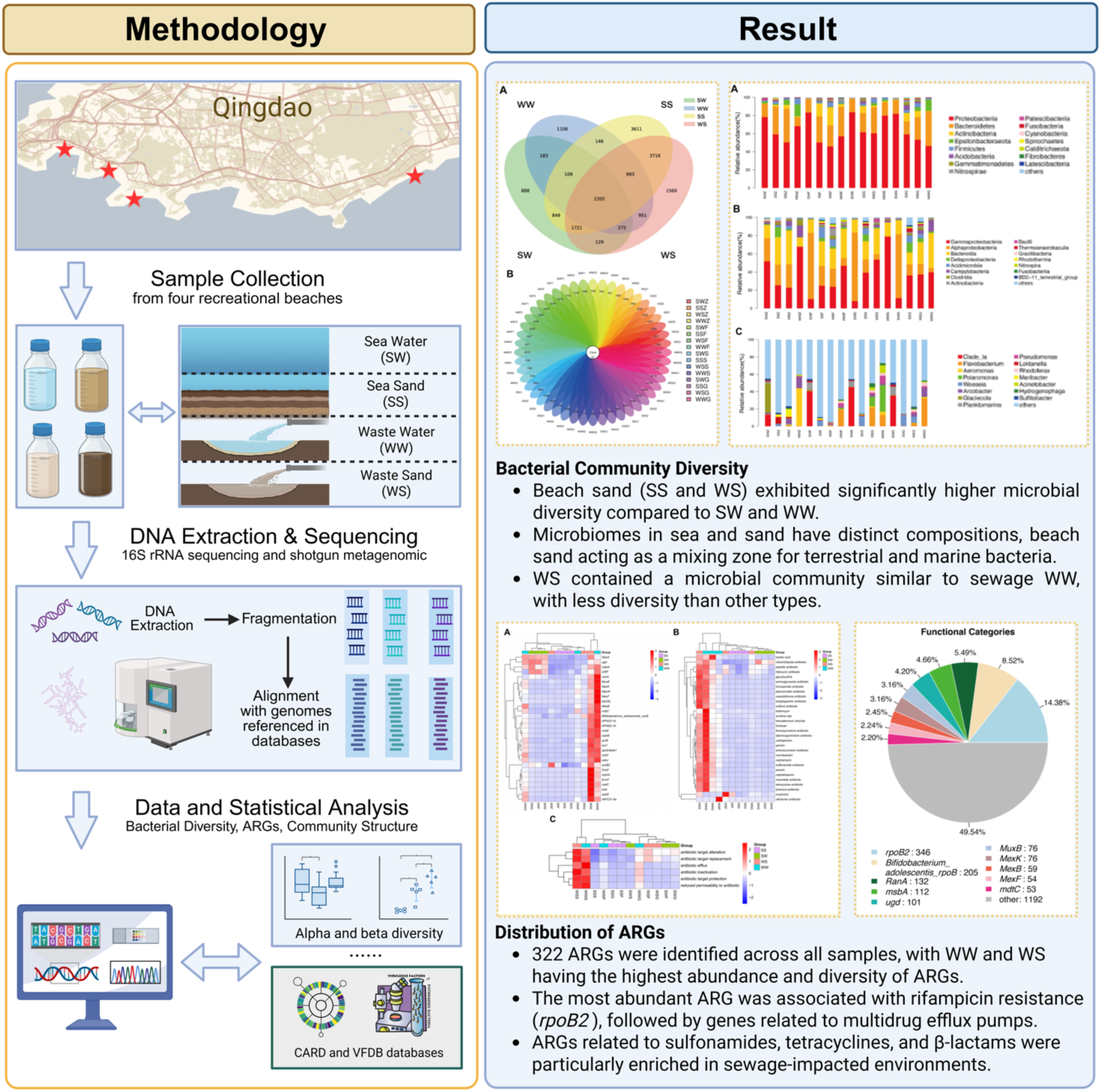

## 1. Introduction

Antibiotic resistance has emerged as a pressing global health threat, with drug-resistant infections claiming an estimated 700,000 lives each year. The widespread use and misuse of antibiotics in human and veterinary medicine have led to the proliferation of antibiotic-resistant bacteria and antibiotic resistance genes (ARGs) in various environments ^1–3^. Previous studies have identified municipal wastewater, riverine systems, and sediments as important hotspots for the dissemination of ARGs into the environment ^4–7^. Coastal beach sands, which filter and accumulate organic and inorganic matter from both land and sea, provide a multitude of microbial niches ^8,9^. In popular recreational beaches, these environmental inputs are compounded by intense human activity, potentially leading to dynamic and unique bacterial communities.

Fecal indicator bacteria, such as *Enterococcus* or *Escherichia coli*, are routinely monitored in beach water and sand as proxies for sewage contamination; their presence can reflect environmental quality and is directly tied to the health and safety of beach users, as well as the economic prosperity of coastal communities ^10,11^. However, beyond standard fecal indicator bacteria counts, relatively few studies have investigated the broader spectrum of pathogenic bacteria and ARGs in beach environments ^12^. Recreational beaches with heavy tourist traffic merit special attention because prolonged or frequent exposure to sand and water enriched with ARGs or pathogens could pose significant health risks to the public ^13,14^.

ARGs introduced via wastewater or runoff can persist and spread in coastal ecosystems, potentially creating a reservoir that facilitates the movement of resistance traits between environments. The discharge of untreated or partially treated sewage, especially during storm events or sewer overflows, has been shown to alter beach microbiomes and introduce human-associated bacteria ^15,16^. Likewise, hospital effluents and urban runoff have been implicated in disseminating ARGs into natural waters ^5^. Because water can readily transport microorganisms and genetic material, it is crucial to track how antibiotic-resistant bacteria and ARGs move through connected environments and to develop strategies to mitigate this spread ^17,18^.

In this study, we investigated the distribution and diversity of bacterial communities and ARGs at four geographically distinct but high-use beaches in Qingdao, China. By analyzing multiple sample types, including seawater (Sea Water, SW), beach sand from the intertidal zone (Sea Sand, SS), sand adjacent to sewage outlets (Waste Sand, WS), and the sewage water discharges themselves (Waste Water, WW), we aimed to unravel the extent to which sewage inputs and seawater influence the beach sand microbiome and resistome. High-throughput 16S rRNA amplicon sequencing and shotgun metagenomic sequencing were employed to profile microbial taxa, ARGs, and virulence factors. We hypothesized that beach sands serve as a convergence point for microbes from land (sewage) and sea, potentially accumulating a diverse array of ARGs and opportunistic pathogens. Our results highlight that beach sand is indeed an under-recognized reservoir of antibiotic resistance and virulence genes. This work underlines the need to include beaches in routine environmental surveillance for antimicrobial resistance and informs management practices to reduce resistome-related health risks in coastal recreational areas.

## 2. Material and Methods

### 2.1 Study site and sample collection

This study was conducted at four coastal sites in Qingdao, Shandong Province, China: Zhan Qiao (Z), First Bathing Beach (F), Second Bathing Beach (S), and Sculpture Garden (G). All four locations are popular recreational beaches that differ in geomorphology and receive large numbers of visitors. At each site, we collected four types of samples to represent different environmental compartments: Sea Water (SW) from the nearshore surface ocean, Sea Sand (SS) from the intertidal and subtidal beach zones, Waste Sand (WS) from sand in the supratidal zone near sewage or stormwater discharge points, and Waste Water (WW) representing the discharge (sewage or runoff) entering the beach. All samples were collected on March 15, 2021, during low tide at midday, with sampling at each site completed within approximately a one-hour period to minimize temporal variability. Triplicate samples were taken for each sample type at each site. Water samples of approximately 5 liters each were collected in sterile plastic containers and transported to the laboratory on ice, where they were processed within 48 hours. Sand samples weighing between 500 and 1000 grams each were collected using sterile plastic bags and were immediately stored at -80 °C until DNA extraction.

**Figure 1.**
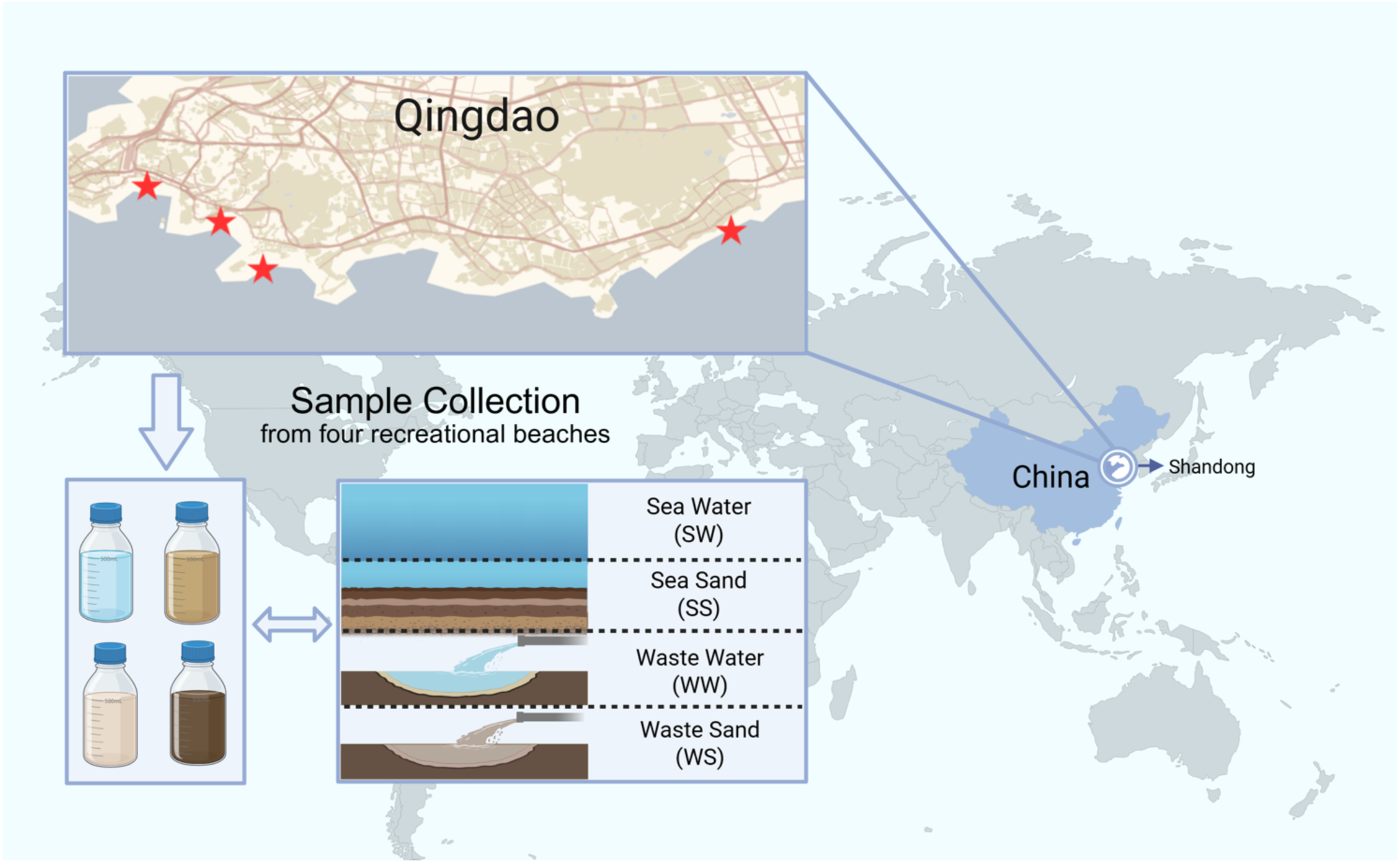
Map of sampling sites in Qingdao, China, where samples were collected. The four recreational beach sites, Zhan Qiao, First Bathing Beach, Second Bathing Beach and Sculpture Garden, are indicated on the map. At each site, SW samples were taken from nearshore waters, SS from intertidal sand, WS from beach sand adjacent to sewage outfalls, and WW from the effluent discharge flowing onto the beach.

### 2.2 DNA Extraction and 16S rRNA Amplicon Sequencing

Total genomic DNA was extracted from each sample using a commercial DNA extraction kit. DNA concentration and purity were checked with a NanoDrop spectrophotometer and by agarose gel electrophoresis. For bacterial community profiling, the V3-V4 hypervariable regions of the 16S rRNA gene were amplified by PCR using the universal primers 343F (5’TACGGRAGGCAGCAG-3’) and 798R (5’-AGGGTATCTAATCCT-3’). PCR reactions (in triplicate for each sample) were carried out with Tks Gflex DNA Polymerase (Takara) and appropriate barcoded adapters for downstream sequencing. Amplicon size and quality were verified by 1% agarose gel electrophoresis. PCR products were purified using AMPure XP beads (Agencourt), then subjected to a second round of PCR to attach Illumina sequencing indices and adapters. The resulting amplicon libraries were purified again with AMPure XP beads and quantified using a Qubit dsDNA HS Assay kit (Thermo Fisher Scientific). Equimolar amounts of each indexed library were pooled, and the pooled library was sequenced on an Illumina platform (paired-end mode) to generate 16S rRNA amplicon reads.

### 2.3 Shotgun Metagenomic Sequencing

For metagenomic analysis, shotgun libraries were prepared from the same DNA extracts representing each sample. Approximately 100 ng of DNA per sample was used to construct Illumina paired-end libraries with an average insert size of about 350 bp, following standard protocols. Libraries were sequenced on an Illumina high-throughput sequencer to produce paired-end reads (150 bp read length).

### 2.4 Amplicon Sequence Processing and OTU Analysis

Raw 16S rRNA amplicon reads in FASTQ format were quality-filtered using Trimmomatic v0.36 ^19^ to remove Illumina adapter fragments, trim low-quality regions (using a sliding window of 4 bases with average quality < Q20), and discard reads shorter than 200 bp. Forward and reverse reads passing filter were merged into full-length amplicon sequences with FLASH v1.2.11 ^20,21^ using a minimum overlap of 10 bp and maximum overlap of 200 bp (mismatch rate ≤ 20%). Further denoising was performed by removing reads containing ambiguous bases or homopolymer runs and by filtering out chimeric sequences using the UCHIME algorithm in QIIME v1.8.0 ^22^. The high-quality reads were then clustered into operational taxonomic units (OTUs) at 97% sequence similarity using VSEARCH ^23^. The most abundant sequence from each OTU cluster was selected as the representative sequence. Taxonomic classification for each representative OTU was assigned using the RDP Classifier ^24^ against the SILVA 16S rRNA database (version 132), with a confidence cutoff of 70%. OTU abundance tables were generated for downstream community analyses.

#### Metagenomic Assembly and Gene Annotation

Quality control of raw metagenomic reads was performed with Trimmomatic ^19^ using similar parameters to the 16S pipeline with adapters and low-quality sequences removed. Any reads identified as potential host contamination such as human DNA were removed by mapping against the human reference genome (hg19) using Bowtie2 v2.2.9 ^25^, and mapped reads were discarded. The remaining high-quality reads from each sample were de novo assembled into contigs using MEGAHIT v1.1.2 ^26^ with default settings. Resulting scaffolds were broken at ambiguous regions to generate contigs (scaftigs). Contigs shorter than 500 bp were discarded to focus on reliably assembled sequences. Gene prediction was carried out on the assembled contigs using Prodigal v2.6.3 ^27^ in metagenomic mode, identifying open reading frames (ORFs) and translating them into amino acid sequences. All predicted proteins from all samples were pooled and clustered to construct a non-redundant gene catalog. CD-HIT v4.6.7 ^28^ was used to cluster sequences at 95% amino acid identity and 90% coverage of the longer sequence. For each cluster, the longest representative sequence was retained as the reference gene. To quantify gene abundance, we mapped quality-filtered reads from each sample back to the non-redundant gene set using Bowtie2 (95% identity threshold). The number of reads mapping to each gene in each sample was recorded, and gene abundance was normalized for each sample to account for variability in sequencing depth. For functional annotation, representative gene sequences were searched against several databases using BLASTp ^29^ with an e-value cutoff of 10^-5^. We used the NCBI non-redundant protein database (NR) for broad taxonomic identification, as well as KEGG (Kyoto Encyclopedia of Genes and Genomes) for pathway annotation ^30,31^, the Clusters of Orthologous Groups (COG) database for functional categories, and the Swiss-Prot/UniProt database for high-quality protein function annotation. Gene Ontology (GO) terms were also assigned where possible. Taxonomic assignments for each gene were derived from the best NR hits and were used to profile the community composition at the domain, phylum, class, and genus levels by summing gene abundances belonging to the same taxa.

### 2.5 Identification of Antibiotic Resistance and Virulence Genes

To characterize the resistome in our samples, the assembled gene catalog was compared against the Comprehensive Antibiotic Resistance Database (CARD, version 3.0.9) ^32^ using BLASTp. Genes with significant matches to known ARGs in CARD were identified, and each gene was annotated with its corresponding resistance ontology (antibiotic class, resistance mechanism, and gene family). We calculated the relative abundance of each ARG by summing the abundances of all genes matching that ARG or gene family in a given sample. Similarly, to screen for potential virulence factors, we aligned the gene catalog against the Virulence Factor Database (VFDB) ^33^. Identified virulence genes were catalogued for each sample type.

### 2.6 Statistical and Diversity Analyses

Alpha-diversity indices such as Chao1 richness, Shannon index, and Simpson index were calculated for each sample based on the 16S OTU table. Differences in diversity between sample types were visualized using violin plots. Beta-diversity patterns were examined through ordination and clustering analyses. Principal Coordinates Analysis (PCoA) was performed on Bray-Curtis dissimilarity matrices of OTU relative abundances to visualize differences in community structure among samples. We also conducted hierarchical clustering of samples using the unweighted pair-group method with arithmetic mean (UPGMA) based on OTU composition similarity. All statistical analyses and visualizations were carried out in R (v3.6.0) unless otherwise noted. Heatmaps to compare ARG composition across samples were generated using the pheatmap package in R, and Venn diagrams for OTU overlap were created with the VennDiagram package.

## 3. Results

### 3.1 General characteristics of samples

#### 3.1 OTU Richness and Overlap Among Sample Types

Across all samples and sites, a total of 18,448 bacterial OTUs were identified from the 16S rRNA amplicon data, based on clustering at 97% sequence similarity (Supplementary Table 1). A subset of 2,202 OTUs approximately 12% of the total was shared among all four sample types (SW, SS, WS, WW), representing the core microbiome present in every environment. Beyond this core, each sample type harbored a considerable number of unique OTUs. For instance, SS samples contained the largest pool of unique OTUs compared to other sample types, whereas WW and SW samples had the fewest unique OTUs. Notably, sand samples consistently exhibited higher OTU richness than water samples at all sites. In other words, both SS and WS had a greater number of observed OTUs than the corresponding SW or WW from the same location. These results suggest that beach sands, particularly the intertidal sand, support a more diverse bacterial assemblage than the beach water or sewage effluent.

**Figure 2.**
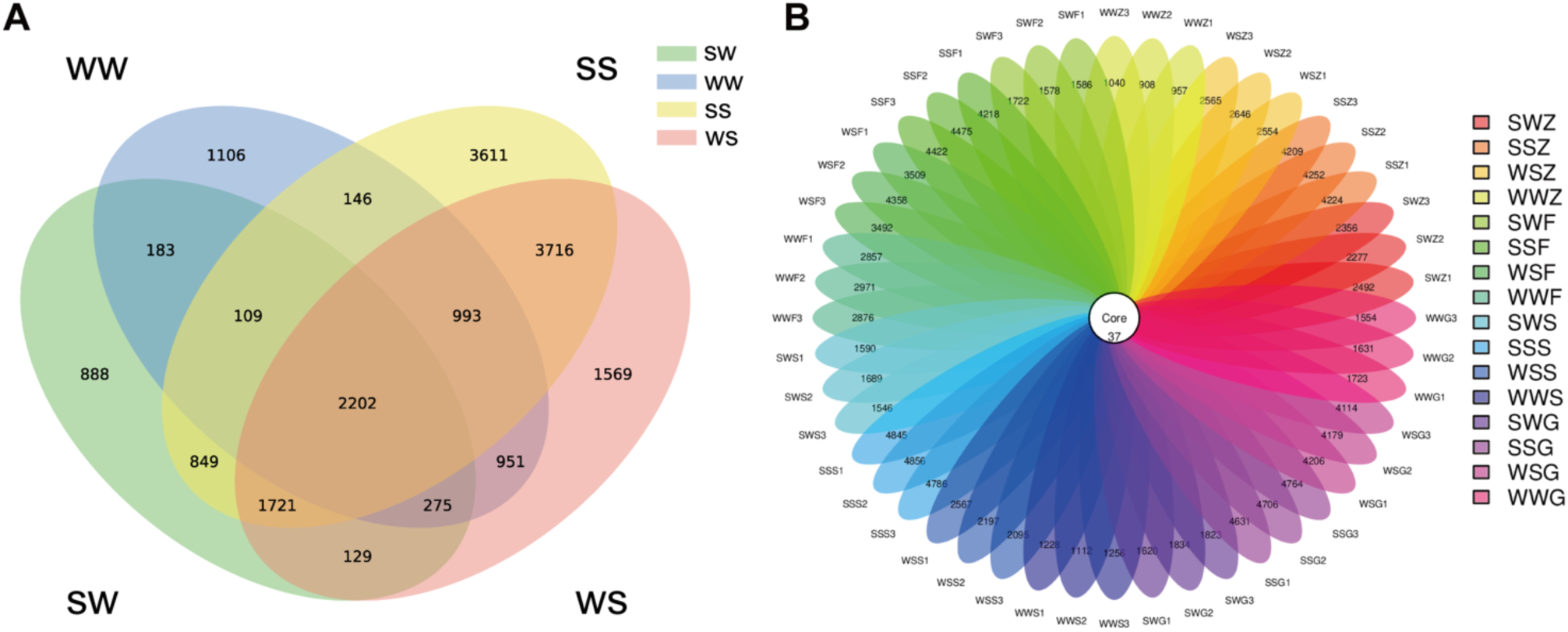
OTU richness and overlap among sample types. (A) Venn diagram showing the overlap of OTUs detected in sea water (SW, green), waste water (WW, blue), sea sand (SS, yellow), and waste sand (WS, red) samples. The number in the center represents OTUs common to all sample types, referred to as core OTUs, while numbers in each ellipse indicate OTUs unique to that particular sample type or shared only with a subset of others. (B) “Flower” plot showing the total OTU count for each sample type across all sites. The center of the flower corresponds to the core OTUs, meaning those present in all sample types, and each petal represents the OTUs unique to one sample type after excluding the core overlap. Sampling locations include Zhan Qiao (Z), First Bathing Beach (F), Second Bathing Beach (S), and Sculpture Garden (G).

### 3.2 Bacterial Community Composition

The bacterial community profiles differed notably among seawater, sand, and sewage-associated samples, although a few major taxa dominated across all samples. At the phylum level, Proteobacteria and Bacteroidetes were the most abundant phyla in every sample, together accounting for over 86% of the sequence reads on average (Figure 3A).

**Figure 3.**
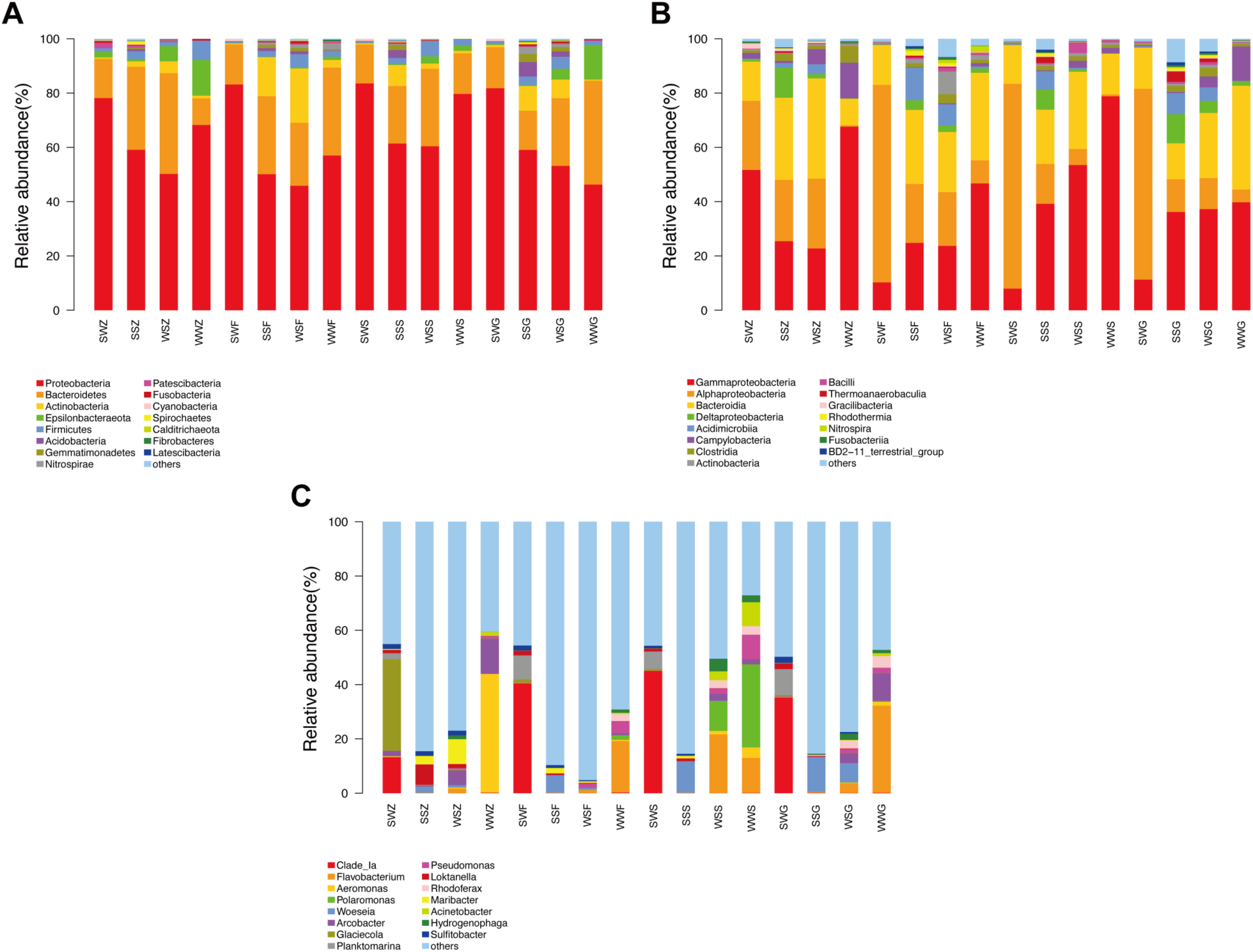
Taxonomic composition of bacterial communities in different sample types. Stacked bar charts show the relative abundance of dominant taxa in each sample (each bar represents a single sample). (A) Phylum-level composition: Proteobacteria and Bacteroidetes dominate across all samples, with noticeable contributions of Actinobacteria in sand samples and Epsilonbacteraeota and Firmicutes in sewage-associated samples. (B) Class-level composition: Gammaproteobacteria, Alphaproteobacteria, and Bacteroidia are the top classes. Gammaproteobacteria are particularly enriched in WW samples, whereas Alphaproteobacteria dominate SW samples. (C) Genus-level composition: Only the most abundant genera are colored; less abundant taxa are grouped as “Other”. The genera composition varies by sample type. For example, SAR11 Clade Ia and *Planktomarina* are prominent in SW, *Flavobacterium* is notable in WW and WS, *Pseudomonas* is elevated in WS, and *Woeseia* dominates many SS samples.

In addition to these, Actinobacteria was a prominent phylum in the sand samples (both SS and WS), whereas Epsilonbacteraeota and Firmicutes were relatively enriched in the WW and WS samples compared to the seawater and clean sand. These patterns indicate that sewage-impacted matrices contained bacterial groups less common in open seawater. At the class level, the communities were largely composed of Gammaproteobacteria, Alphaproteobacteria, and Bacteroidia, which together constituted roughly 80-85% of the bacteria in most samples (Figure 3B). Gammaproteobacteria was the single most dominant class in all WW samples, reflecting the high load of sewage-derived bacteria such as Enterobacteriaceae and other gammaproteobacterial groups in the effluents. In contrast, Alphaproteobacteria, which include many marine oligotrophic bacteria like the SAR11 clade, were relatively more abundant in the SW samples. The sand samples (SS and WS) showed mixed dominance of both Gammaproteobacteria and Alphaproteobacteria, along with Bacteroidia. Minor classes such as Deltaproteobacteria and Acidimicrobiia were also detected at lower abundances. At the genus level, we observed substantial variation in community composition between sample types and sites. Many of the most abundant OTUs could only be classified to higher taxonomic levels, indicated as “unclassified” or family-level taxa, suggesting a large proportion of novel or not well-characterized bacteria. Nonetheless, among the identifiable genera, we found clear habitat preferences. In the SW samples, marine heterotrophs like the SAR11 clade (Clade Ia) and *Planktomarina* (family Rhodobacteraceae) were dominant. In the WS and WW samples, genera associated with organic pollution were prevalent: for example, *Flavobacterium* (a genus of Bacteroidetes often found in nutrient-rich waters) was highly represented in both WS and WW. The genus *Pseudomonas*, known for its metabolic versatility and presence in soil and water, including contaminated sites, formed a relatively large fraction of the WS communities. Meanwhile, the SS samples were often dominated by the genus *Woeseia* (family Woeseiaceae, order Chromatiales), a group of Gammaproteobacteria commonly found in marine sediments. These genus-level trends indicate that the microbial communities in beach sands reflect a combination of marine and terrestrial (pollution-derived) influences, with sewage-associated bacteria thriving in the waste-affected sand and more typical marine bacteria in the seawater and clean sand.

### 3.3 Alpha and Beta Diversity of Communities

We evaluated bacterial diversity within each sample (alpha-diversity) and between samples (beta-diversity) to further resolve differences among the four sample types. Alpha-diversity indices calculated from the OTU data (Supplementary Table 2) showed clear trends: beach sand contained a more diverse bacterial assemblage than seawater or wastewater. In particular, the Simpson diversity index was highest in SS samples, followed by WS, then WW, and was lowest in SW samples (Figure 4A). The Shannon index displayed a similar pattern (Figure 4B), with SS samples having nearly double the diversity of SW in some cases. These metrics confirm that the heterogeneous sand environment supports greater bacterial diversity, while the open water, especially seawater, has comparatively lower diversity. Interestingly, the seawater sample from Zhan Qiao had a slightly elevated diversity relative to seawater at the other sites, but the overall trend of SS > WS > WW > SW held true across all locations. Differences in community composition between samples (beta-diversity) were visualized via PCoA ordination. When examining each site separately, the four sample types formed distinct clusters in ordination space (Figure 5A-D). In all cases, the triplicate SW samples clustered tightly together and were clearly separated from the clusters of sand and wastewater samples along the principal coordinate axes. For example, at each beach site, the SW cluster was positioned far from the SS cluster, indicating that seawater communities were compositionally distinct from those in beach sand. WS and WW samples at a given site tended to plot closer to each other in the PCoA, reflecting their shared influence from sewage. These patterns suggest that within any given beach, the microbiome of the seawater is quite different from that of the sand, whereas the sand near sewage outfalls (WS) is more similar to the sewage water (WW) source. We further confirmed these relationships by hierarchical clustering of samples using UPGMA (Figure 6A–D). The UPGMA dendrograms grouped samples primarily by their type rather than by site: all SW samples across different beaches clustered together, as did all SS samples, and so on. This indicates that the four sample categories each have characteristic community structures regardless of location. Notably, in both PCoA and UPGMA analyses, SW was the most distinct type, and its samples did not cluster with any non-SW sample, underscoring that the open coastal water harbored a unique subset of the bacterial community not strongly overlapping with the beach sand or sewage. These diversity analyses demonstrate that beach sand microbiomes are richer and, in composition, more similar to sewage-associated communities than to seawater communities. The constant washing of sand by waves and tides does not homogenize the sand and water microbiomes; instead, sand appears to retain a distinct and more complex community.

**Figure 4.**
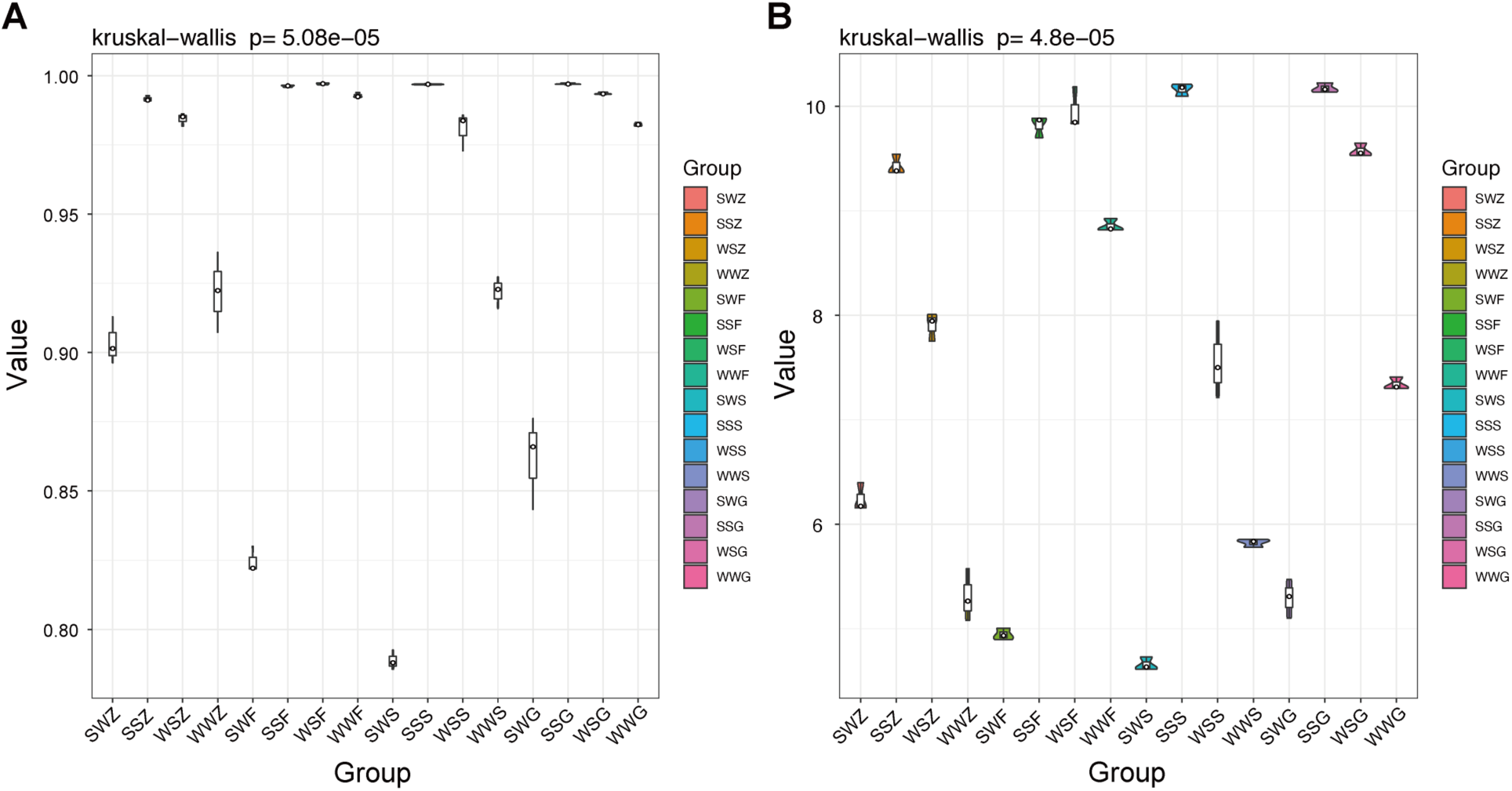
Alpha-diversity of bacterial communities in different sample types. Violin plots summarize the distribution of diversity index values for each sample type (pooling data from all sites). (A) Simpson diversity index, which gives more weight to abundant species, is highest on average in SS samples and lowest in SW, indicating greater evenness and richness in sand communities. (B) Shannon index, which accounts for both richness and evenness, shows a similar trend of SS > WS > WW > SW. The width of each violin represents the density of samples with a given diversity value, and the median is indicated inside each violin.

**Figure 5.**
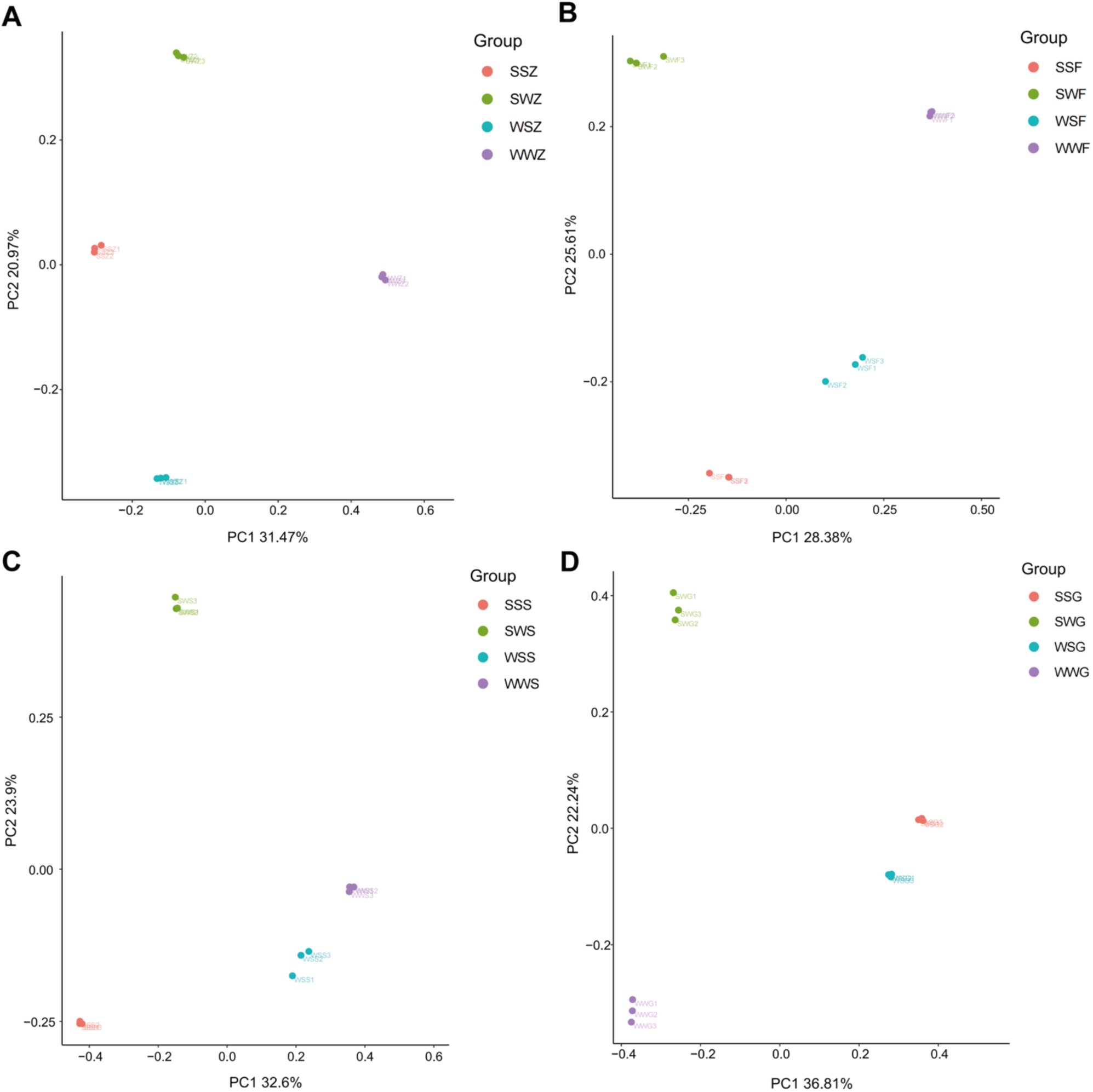
Principal Coordinates Analysis (PCoA) of community composition at each site. PCoA ordinations were performed based on Bray-Curtis dissimilarities of OTU relative abundances. Each panel corresponds to one beach site: (A) Zhan Qiao, (B) First Bathing Beach, (C) Second Bathing Beach, and (D) Sculpture Garden. Within each panel, symbols represent samples (colored by type: green for SW, red for SS, blue for WS, purple for WW). Triplicate samples of the same type cluster together.

**Figure 6.**
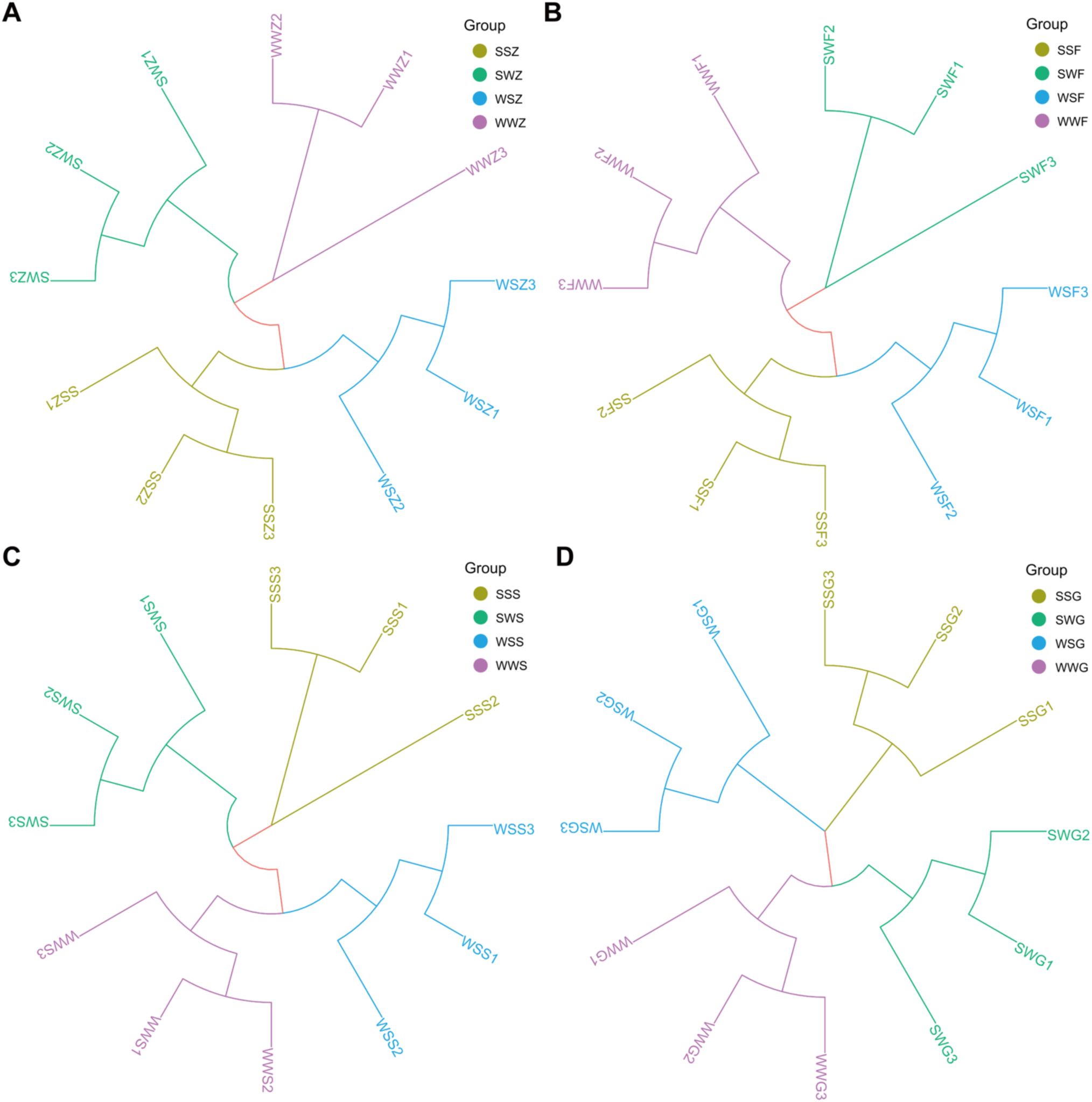
Hierarchical clustering (UPGMA) of samples based on bacterial community similarity. Dendrograms depict the clustering of samples (using Bray-Curtis distance on OTU profiles) for each site: (A) Zhan Qiao, (B) First Bathing Beach, (C) Second Bathing Beach, (D) Sculpture Garden. Samples are color-coded by typeA, green for SW, yellow for SS, blue for WS, purple for WW. In each case, samples clearly cluster by type, reinforcing that each environment harbors a distinct bacterial community.

### 3.4 Distribution of Antibiotic Resistance Genes

Shotgun metagenomic sequencing allowed us to characterize the repertoire of ARGs present in each sample type. In total, we identified 322 distinct ARGs across all samples based on CARD annotations. These resistance genes spanned 33 different antibiotic classes and 6 major resistance mechanisms (Supplementary Table 3 and Supplementary Table 4). The ARGs detected include those conferring resistance to nearly all major antibiotic families, indicating a diverse resistome in this coastal environment. The most frequently detected resistance gene by cumulative abundance across samples was *rpoB2*, a variant of the RNA polymerase β-subunit gene associated with rifampicin resistance (Figure 7; Supplementary Table 7). In terms of prevalence by antibiotic class, genes conferring resistance to peptide antibiotics, such as bacitracin and polymyxins, were among the most abundant in our dataset, and multidrug efflux pump genes were also ubiquitous. Not surprisingly, the highest richness and abundance of ARGs were observed in the sewage-contaminated samples. On average, a single WW sample contained approximately 86% of all the distinct ARG types catalogued in this study, indicating that nearly every resistance gene identified across all samples was also present in WW. This highlights raw sewage as a major reservoir of diverse ARGs. WS samples, which are directly impacted by the sewage outfalls, carried on average about 72% of the total ARG types. By contrast, the clean beach sand (SS) samples contained on average only around 47% of the ARG types, and SW samples about 60%. Thus, relative to WW and WS, the SW and SS resistomes were less diverse. In addition to diversity, the overall load of ARGs (cumulative abundance) was much higher in WW and WS than in SS and SW.

**Figure 7.**
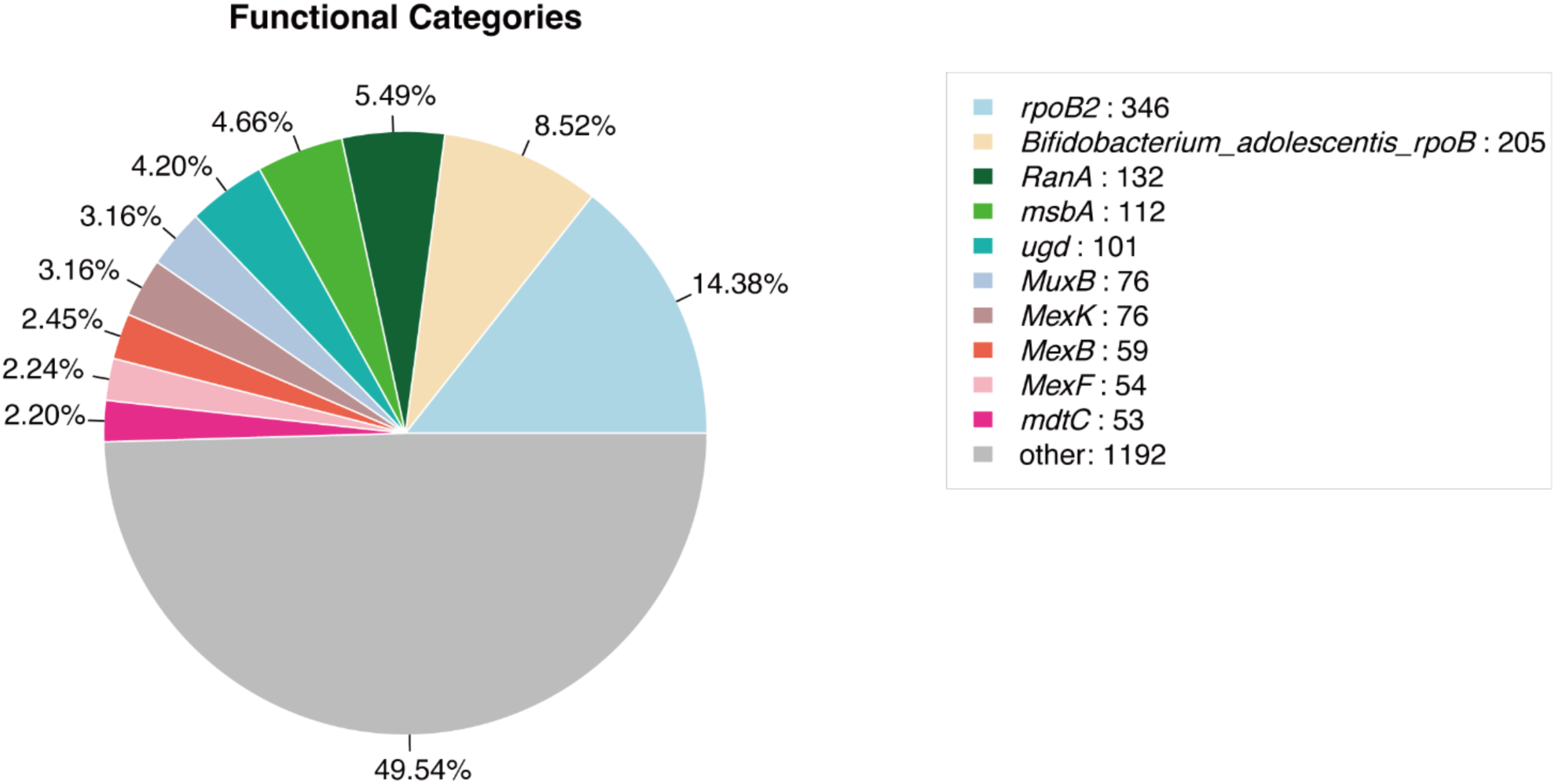
Top 10 most abundant ARGs identified across all samples. Bar chart ranks ARGs by their total normalized abundance across all samples, based on the sum of gene abundances. The rifampicin resistance gene *rpoB2* is the single most abundant ARG in the dataset, followed by various multidrug efflux pump genes and other resistance determinants. The prominence of *rpoB2*, a gene typically associated with high-level rifamycin resistance in bacteria, highlights the strong selective pressures present in these environments. Each ARG bar is colored by its resistance mechanism or drug class for visual distinction.

Figure 8 illustrates this clearly: The heatmap of the top 30 individual ARGs (Figure 8A) revealed several highly abundant ARGs consistently enriched in sewage-related samples. Prominent among these were genes conferring resistance to sulfonamides (*sul1*, *sul2*), tetracyclines (*tetM, tetX*), macrolides (*ermB*), and β-lactams (including *bla* variants). The ubiquitous presence and high intensity of *sul1* and *sul2*, particularly in WS samples, align with their frequent association with integrons and multidrug resistance elements widely reported in wastewater-influenced environments. Similarly, *tetM* and *tetX* genes were highly abundant, reflecting historical and ongoing tetracycline use. Genes such as *ermB*, encoding ribosomal modification enzymes, and β-lactamase variants were also prominent, indicating resistance to clinically important macrolides and β-lactam antibiotics. These ARGs’ higher abundance in WS compared to SS or SW highlights the concentration of ARGs in sludge-like matrices near sewage outlets, with markedly reduced levels in seawater, indicative of dilution and natural attenuation processes.

**Figure 8.**
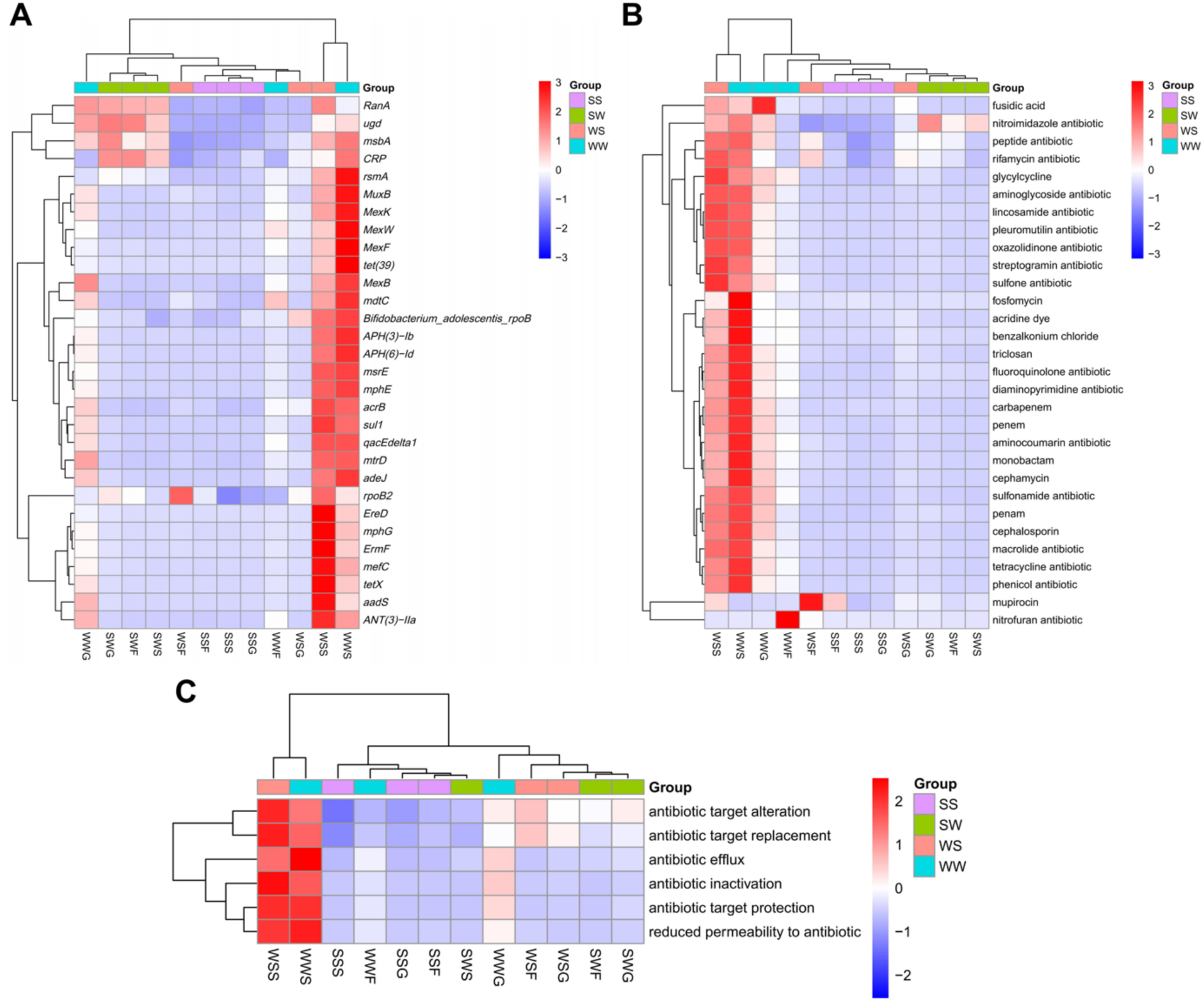
Distribution and diversity of ARGs across sample types. Heatmaps illustrate the relative abundances of ARGs grouped by (A) individual gene (ARO_Name), (B) drug class, and (C) resistance mechanism in SW, SS, WS, and WW samples. Each row represents one of the 30 most abundant ARGs or categories in that grouping. Color intensity from blue (low) to red (high) on a log scale reflects relative abundance across the four sample types.

Analyzing ARGs by antibiotic class (Figure 8B) underscored the dominance of tetracycline, sulfonamide, β-lactam, and macrolide resistance across the samples. Tetracycline and sulfonamide resistance classes exhibited the highest overall abundance, driven by genes such as *tetM/X* and *sul1/2*, mirroring the gene-level data. β-lactam resistance, including resistance to extended-spectrum cephalosporins, also showed substantial enrichment, especially in WS. Additionally, macrolide resistance presented notable abundance, again reflecting sewage influence. Importantly, lower yet detectable levels of resistance genes to clinically critical antibiotics like aminoglycosides, quinolones, and polymyxins were observed, suggesting potential health risks associated with ARG dissemination from wastewater into recreational environments.

Further categorization based on resistance mechanisms (Figure 8C) revealed efflux pumps and antibiotic-inactivating enzymes as the dominant resistance strategies across all environments, particularly in sewage-influenced matrices (WS and WW). Efflux pump genes, known for conferring broad-spectrum multidrug resistance, were the most widespread and abundant, reflecting their role in bacterial adaptation to diverse antimicrobial stressors. Enzymatic inactivation, exemplified by β-lactamases and aminoglycoside-modifying enzymes, was also highly prevalent, especially in WS. Target protection and modification mechanisms (e.g., ribosomal protection by *tetM*, rRNA methylation by *ermB*) similarly featured prominently. The diverse array of resistance mechanisms in sewage-associated samples underscores their role as hotspots for ARG evolution and horizontal gene transfer, potentially facilitating ARG dissemination into downstream ecosystems.

### 3.5 Occurrence of Bacterial Virulence Factors

In addition to ARGs, we screened the metagenomes for virulence factor genes (VFs) to assess the presence of potential pathogens or pathogenic traits in these microbial communities. A wide array of VFs was detected, including genes related to adhesion, toxin production, secretion systems, iron acquisition, and other pathogenic mechanisms based on VFDB matches. Interestingly, the occurrence and diversity of virulence factors were highest in the beach sand samples, particularly in SS. The SS samples contained a greater variety of distinct VF genes compared to WW, WS, or SW samples. Many of these virulence genes are associated with clinically relevant pathogens. For example, genes for hemolysins, type III secretion system components, and quorum sensing regulators were observed. The elevated VF diversity in sand could be due to sand serving as a long-term sink for pathogens from various sources such as human, animal, and environmental, which then persist or proliferate in the sand matrix. By contrast, although WW introduces many pathogens, its bacterial community may be dominated by a few highly abundant strains, thereby limiting the apparent diversity, and SW may not allow long-term persistence of certain pathogens due to environmental stresses. These results point to beach sands as not only reservoirs of ARGs but also as reservoirs of virulence genes, which could potentially be transferred among bacteria in the sand environment or picked up by humans through contact.

### 3.6 Functional Gene Profiles

To gain a broader understanding of the functional potential of the beach microbiome, we annotated metagenomic genes against metabolic and cellular pathways in KEGG. The microbial communities across our samples showed enrichment in genes for core metabolic functions that are fundamental in marine and terrestrial ecosystems. The ten most abundant functional categories at KEGG Level 3 pathway detail included pathways and systems such as ABC transporters, Ribosome, Aminoacyl-tRNA biosynthesis, Two-component regulatory systems, Purine metabolism, Porphyrin and chlorophyll metabolism, Oxidative phosphorylation, Quorum sensing, RNA degradation, and RNA polymerase (Figure 9). These highly abundant functions indicate that the microbiota in beach sand and water are actively involved in nutrient transport, translation and transcription processes, energy production and intercellular communication, reflecting a community well-equipped for environmental survival and nutrient cycling. At a broader level, based on KEGG Level 1 and 2 classifications, genes associated with Metabolism were the most dominant, particularly those involved in carbohydrate metabolism, amino acid metabolism and energy metabolism, which were substantially over-represented compared to other categories. This is consistent with the notion that these communities are degrading organic matter and cycling nutrients in the beach environment (Falkowski et al., 2008; Gobet et al., 2012). Meanwhile, a significant proportion of the annotated genes fell under the Human Diseases category of KEGG. Within this category, pathways related to infectious diseases and drug resistance were prominent, such as genes for microbial adherence, invasion, toxin production, as well as bacterial defense against antibiotics. There were also genes mapping to pathways implicated in cancers, which often reflects the presence of certain microbial genes involved in xenobiotic metabolism or in the modulation of host cellular processes, rather than the actual occurrence of disease. The prevalence of “human disease” related genes align with our direct detection of ARGs and VFs, reinforcing that these beach and sewage-associated microbiomes harbor a pool of functions relevant to pathogenicity and resistance.

**Figure 9.**
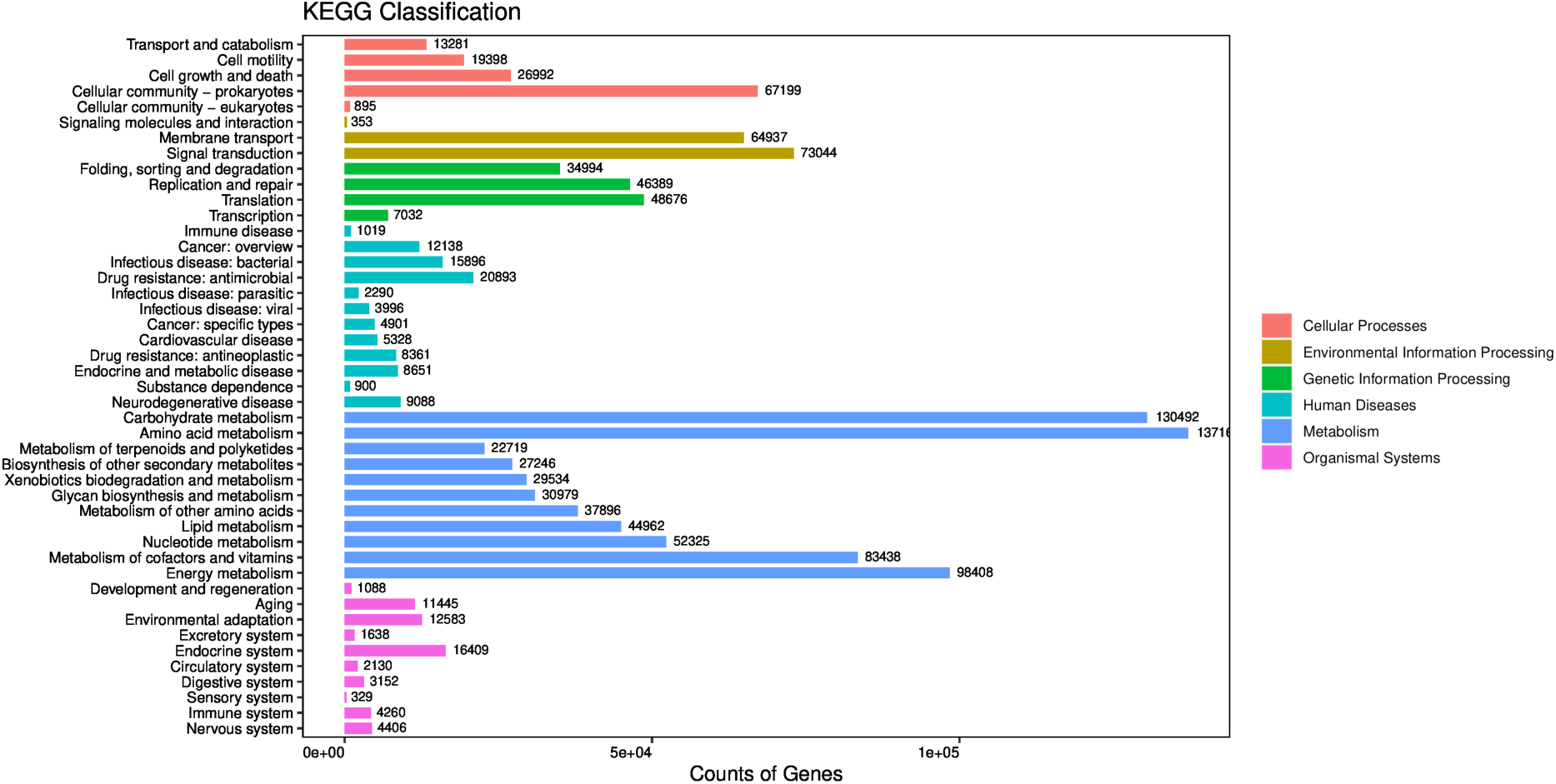
KEGG functional annotation summary. Bar charts show the number of genes assigned to major KEGG functional categories in the combined gene catalog. (A) Distribution of genes across KEGG Level 1 categories (e.g., Metabolism, Environmental Information Processing, Genetic Information Processing, Cellular Processes, Human Diseases). Metabolism-related genes constitute the largest share, but notably, a considerable number of genes fall into the Human Diseases category as well. (B) Distribution of genes within selected Level 2/3 pathways. The most enriched pathways include basic cellular machinery and metabolic pathways (e.g., ribosomal proteins, ABC transporters, etc., highlighted in text). Additionally, genes associated with infectious disease pathways and antibiotic resistance mechanisms are well represented, consistent with the resistome and virulome analysis.

## 4. Discussion

Previous research on environmental antibiotic resistance has largely focused on habitats such as wastewater treatment plants, rivers, estuaries, and marine water columns ^34–38^. Beaches, especially the sand, have received comparatively less attention in terms of microbial ecology and resistome characterization, aside from studies examining specific pollution events ^15^ or monitoring fecal indicator bacteria ^10,11,39^. In this study, we simultaneously surveyed the microbiomes of beach sand, seawater, and sewage inputs at recreational beaches, providing a holistic view of how these interconnected compartments influence each other. Our results clearly show that beach sands are not microbiologically isolated or benign environments; rather, they are dynamic reservoirs shaped by both terrestrial (sewage) and marine influences and capable of harboring diverse bacteria, including potential pathogens and ARGs. This has important implications for public health, as it expands our understanding of where antibiotic resistance may be developing and persisting outside of clinical settings.

### 4.1 Bacterial Community Composition Reflects Anthropogenic Impact

The taxonomic analysis revealed that Proteobacteria and Bacteroidetes dominate the beach sand and water microbiomes, which is in line with many other marine and coastal microbiome studies. Bacteroidetes are particularly noteworthy as they include many gut-associated bacteria, implying a degree of fecal contamination in these environments ^40–42^. The co-dominance of Proteobacteria, one of the largest bacterial phyla encompassing a wide variety of metabolisms, indicates a mix of autochthonous marine bacteria and allochthonous bacteria. Many well-known pathogens fall within Proteobacteria, including *Escherichia coli*, *Salmonella*, *Vibrio cholerae*, *Helicobacter pylori*, so a high proportion of Proteobacteria in these samples could point to the presence of potentially pathogenic species ^43,44^. We also observed Actinobacteria as a significant component of the sand bacterial communities; this phylum includes some environmental species that are medically important such *Mycobacterium* and *Nocardia* spp., which can cause diseases in humans ^45^. The enrichment of Firmicutes in WW and WS (relative to SW and SS) is consistent with known sewage microbiota signatures, as human waste and sewer infrastructure tend to carry many Firmicutes ^46^. At finer taxonomic resolution, the differences between sample types become more indicative of anthropogenic impact. Gammaproteobacteria, a class containing many opportunistic pathogens and pollution-tolerant bacteria, was disproportionately abundant in sewage and waste sand samples. This class includes families like Enterobacteriaceae, Vibrionaceae, and Pseudomonadaceae, which are known to survive across a range of conditions and are often introduced by sewage or runoff ^47^. Alphaproteobacteria, by contrast, were more dominant in open seawater, reflecting typical marine communities such as the SAR11 clade that thrive in oligotrophic conditions ^8,48,49^. The coexistence of these groups in sand, especially WS, suggests that sand is a meeting point for microbes from both sources. We identified several genera of concern that have been noted in other studies of polluted waters. These include enteric bacteria like *Escherichia/Shigella* and *Salmonella*, waterborne bacteria like *Vibrio* and *Arcobacter*, and generalist environmental bacteria like *Pseudomonas* and *Acinetobacter* ^5,35^. All these groups were present to varying degrees in our samples. Importantly, the cumulative relative abundance of bacterial genera that are commonly associated with human infections such as, *Escherichia*, *Salmonella*, *Pseudomonas*, *Acinetobacter*, *Enterococcus*, was highest in WW, followed by WS, then SS, and lowest in SW (Supplementary Table 5). This gradient strongly indicates that sewage discharges concentrate potentially pathogenic bacteria, which then disperse into beach sand with dilution, and only a smaller fraction makes it into the seawater farther away. Thus, while the presence of these bacteria in sand does not mean the sand is causing infections, it does highlight the role of sand as a reservoir for microbes of public health relevance.

### 4.2 Beach Sand as a Hotspot of Bacterial Diversity and Exchange

Our analysis of alpha and beta diversity highlights the complex ecosystem that beach sand represents. Consistent with patterns observed in river sediments versus water ^5,7^, we found that sand (both SS and WS) had higher bacterial diversity than the overlying water. Sand grains offer surfaces for biofilm formation and create microenvironments that can support a wider range of microorganisms than the open water column, which is subject to more fluctuations and generally has lower nutrient concentrations. The finding that SS > WS > WW > SW in diversity suggests that as we move from seawater to pure sewage, diversity decreases, perhaps because extreme environments like raw sewage favor certain hardy taxa and exclude others. The beach sand, sitting at the intersection, seems to gather bacteria from multiple sources, leading to a composite community with high richness. One implication of having these dense, diverse communities in sand is the potential for horizontal gene transfer. The close proximity of various bacteria within sand biofilms can facilitate the exchange of genetic elements, including plasmids and transposons carrying ARGs. Riverine studies have posited that sediments provide a stable habitat where ARGs can spread among bacteria ^5,50,51^. By analogy, the beach sand “microbial reef” may promote interactions that would be rare in open water, such as between marine bacteria and gut bacteria that ended up in sand via sewage. These interactions could allow ARGs and virulence factors to move into new host bacteria. Our results also underscore the influence of sewage-borne bacteria on the beach microbiota. We saw those genera like *Aeromonas*, *Flavobacterium*, *Arcobacter*, *Polaromonas*, *Pseudomonas*, *Rhodoferax*, and *Acinetobacter* were most abundant in WW and showed a stepwise decline in relative abundance from WS to SS to SW (Supplementary Table 6). This points to a clear dispersal gradient: the farther from the sewage source, the lower the contribution of those sewage-associated genera. Similar observations have been made in studies of wastewater effluent plumes in coastal areas ^15,16,52,53^. Wave action and tidal movements certainly mix seawater with sand to some extent, but our PCoA and clustering showed that the seawater community remained distinct. This could be because many sewage-derived bacteria survive better in the nutrient-rich, particulate environment of sand than in open water, or because they physically adhere to sand grains. Indeed, Gobet et al. (2012) noted that sand grains rapidly develop their own biofilm communities, which are not easily washed away by waves. Thus, while the ocean dilutes contaminants and can wash some bacteria away from the beach, a significant portion remains and establishes within the sand. The separation of SW communities from sand communities suggests limited exchange in the other direction as well; that is, the resident sand microbiota, including any introduced through sewage, do not substantially colonize the water column, at least not in a lasting way. However, cells and DNA from sand can still be released into water intermittently. This dynamic interplay among sewage, sand, and sea warrants further study, particularly with finer temporal resolution such as sampling before and after rain events, or over tidal cycles, to see how quickly bacteria from one compartment appear in another.

### 4.3 Beach Resistome Mirrors Sewage Inputs and Environmental Dilution

Our comprehensive ARG analysis indicates that beach sands, especially those near wastewater discharges, accumulate a wide variety of resistance genes. Many of these genes, such as those for multidrug efflux pumps, β-lactamases, and aminoglycoside-modifying enzyme, are unsurprising in a wastewater context and have been reported in sediments receiving wastewater ^6,34,54,55^. The diversity of ARGs found, encompassing 322 distinct genes across 33 classes, reinforces that environmental reservoirs like beach sands can contain a “record” of local antibiotic usage and resistance dissemination. One noteworthy result was the high abundance of *rpoB2* in all sample types. *RpoB2* encodes an altered RNA polymerase β-subunit that confers resistance to rifamycins; its prevalence here could signal that bacteria in these environments have been exposed to rifampicin or other stressors selecting for these mutations. Ekwanzala et al. (2020) similarly found that mutations in housekeeping genes (like *rpoB*) can become prevalent in polluted environments, possibly due to co-selection pressures (heavy metals or other contaminants co-occurring with antibiotics). The detection of *rpoB2* in beach sand underscores that even the sand, often perceived as inert or safe, is subjected to these selective forces and can maintain resistant populations. The stark contrast in ARG richness and abundance between WW/WS and SS/SW highlights sewage as the primary contributor of ARGs. Essentially, the sewage outfalls are injecting a concentrated resistome into the beach ecosystem. Almost every ARG that turned up in seawater or sand was also found in the sewage, indicating that the resistome of the beach is largely a subset of the sewage resistome, minus some losses due perhaps to environmental filtering. The clean sand (SS) had substantially fewer ARG types and lower total ARG abundance, which likely reflects the cleansing effect of the ocean: tidal flushing, UV exposure, and lower nutrient levels in the open beach sand may reduce the survival of many ARG-bearing organisms or genes. Dilution also plays a role: any given volume of sand farther from the outfall receives fewer bacteria from sewage. It is interesting, however, that the seawater (SW) had a higher fraction of ARG types (approximately 60%) than SS (approximately 47%). This could be because seawater samples, while lower in biomass, can transport ARG-bearing bacteria over larger distances, picking up not just local sewage influence but possibly other diffuse sources (such as urban runoff along the coast or boats). Sand, on the other hand, is more localized. In any case, both SS and SW still harbored dozens of ARGs, some of which are clinically relevant. The presence of ARGs in recreational water is a direct exposure risk for swimmers ^17^, and sand can also be an exposure route through skin contact or inadvertent ingestion, especially for children playing on the beach ^56–58^. Our findings strongly support expanding monitoring and management efforts: beach sands should be included in environmental antibiotic resistance surveillance. To protect public health, regulators typically focus on water quality, but our data show that sand can concentrate ARGs and potentially pathogenic bacteria to levels comparable to or higher than the water. Measures such as public advisories after sewage spills, regular sand testing, or even sand remediation might need to be considered in heavily-used beaches.

### 4.4 Functional Potential and Health Implications

The functional gene profiles provide context to the resistome and taxonomic findings. The dominance of core metabolic pathways in all samples indicates that, regardless of source, the microbes are geared towards fundamental life processes such as breaking down organic matter, producing energy, replicating, etc. This is expected for environmental microbiomes and aligns with known roles of coastal microbes in nutrient cycling and ecosystem functioning ^8,59^. In a healthy scenario, these communities contribute to the degradation of pollutants, which may include residual antibiotics and other chemical contaminants, and to the productivity of coastal systems. However, the over-representation of genes in the “Human Diseases” category, includes ARGs and VFs, is a red flag from a health perspective ^60^. Our detection of many virulence factors, especially in sand, suggests that pathogenic bacteria, or at least bacteria carrying pathogenic traits are present in this environment. These could include typical marine pathogens like *Vibrio* spp. (which have known virulence genes for cholera toxin, etc.) or land-derived pathogens like toxigenic *Escherichia coli*. The co-occurrence of ARGs and VFs raises the concern of bacteria that are both pathogenic and drug-resistant. If such bacteria are present on a beach, there is a non-zero chance they could infect a human through a cut, ingestion of sand or water, or other exposure pathways. While our study did not isolate live pathogens or test infectivity, the genetic evidence alone warrants caution. The finding that surfers and swimmers in the UK were colonized by resistant *Escherichia coli* after exposure to coastal water ^17^ provides a real-world corroboration that recreational contact can indeed transfer resistant bacteria to humans. In Qingdao’s beaches, millions of people visit each year; even a very low probability event could become a public health issue when scaled to such large populations. Moreover, beach sand could act as a chronic source as people might not realize it, but repeated visits could expose them to small doses of these microbes. Researchers have advocated for greater examination of environmental reservoirs in the context of emerging pathogens ^7^. Our work adds to that call, demonstrating that a seemingly benign environment like beach sand is teeming with genetic determinants of resistance and virulence. The next step could be to isolate specific strains from sand and water to see if they indeed correspond to known pathogenic species and to test their resistance profiles. Additionally, studying the survival and decay rates of these genes and organisms in sand vs. water would inform risk assessment models. In summary, our findings highlight that recreational beach sands are an intersection of human, environmental, and microbial health concerns. They reflect the impact of anthropogenic pollution by exhibiting shifts in community composition and enrichment of resistomes, while simultaneously fulfilling natural roles in coastal ecology. Managing this delicate balance will be key. For instance, enhancing wastewater treatment to remove not just nutrients but also microbes and ARGs before effluent reaches beaches could mitigate the influx of resistome elements. Creating buffer zones or engineered wetlands between sewage outlets and beach areas might also help filter bacteria. From a policy standpoint, incorporating ARG monitoring in beach safety guidelines could be considered in the future.

## 5. Conclusions

Using a combination of 16S rRNA gene sequencing and shotgun metagenomic analysis, this study provides a comprehensive overview of microbial diversity, ARGs, and virulence factors in seawater, beach sand, and sewage at four popular beaches in Qingdao, China. We found that beach sands harbor diverse and abundant microbial communities, including many bacteria of human health relevance, and that sands adjacent to sewage inputs accumulate a wide array of ARGs, rivaling the resistome of raw sewage. Seawater and sand continuously interact, but sand retains a distinct microbiome that acts as a repository for ARGs and pathogens. These findings confirm that antibiotic resistance is not confined to obvious hotspots like hospitals and wastewater treatment plants but extends into public recreational areas, thus expanding the scope of environments that need attention in the fight against the spread of resistance. Crucially, our results indicate that the microbial composition and resistome of beach sand are strongly influenced by sewage contamination. Beaches can thus serve as sentinels reflecting urban sewage impacts on the coastal environment. The high load of ARGs and virulence genes in sand underlines the importance of including beach habitats in routine surveillance programs for antibiotic resistance and pathogenic microorganisms. From a public health perspective, repeated or prolonged exposure to sand and water enriched in ARGs and potential pathogens could elevate the risk of colonization or infection with antibiotic-resistant bacteria among beachgoers. Based on our findings, we recommend that coastal management strategies aim to reduce direct sewage and runoff discharges onto recreational beaches. Investing in infrastructure to divert or treat wastewater before it reaches beach areas could significantly lower the introduction of ARGs and harmful microbes. Furthermore, it is advisable for environmental authorities to implement routine monitoring of both beach sands and coastal waters for markers of fecal contamination and antibiotic resistance, especially in areas impacted by sewage discharge. By recognizing beach sand as a critical environmental reservoir of antibiotic resistance, we can better protect both ecosystem and human health and implement more effective interventions to curb the dissemination of antibiotic resistance in the natural environment.

## Resource availability

### Lead contact

Further information and reasonable requests for resources and reagents should be directed to and will be fulfilled by the lead contact, Pengfei Cui.

### Materials availability

Requests for materials should be made via the lead contact. All unique/stable reagents generated in this study are available from the lead contact without restriction.

## Data and code availability

- ***Data***: All data reported in this paper will be shared by the lead contact [Pengfei Cui] upon reasonable requests.
- ***Code***: This paper does not report original code.
- ***Additional Information***: Any additional information required to reanalyze the data reported in this article is available from the lead contact upon request.

## Acknowledgments

This work was supported by National Natural Science Foundation of China (No. 31902421).

## Author contributions

Conceptualization, P.C.; Methodology, C.L., H.M. and P.C.; Formal Analysis, C.L., H.M., Y.C., H.G, X.L., J.X. and H.Z.; Investigation, C.L., H.M. and P.C.; Resources, P.C.; Writing - Original Draft, C.L., H.M., Y.C., H.G, and P.C.; Writing - Review & Editing, C.L., H.M., Y.C., H.G, X.L. and J.X.; Funding Acquisition, P.C.; Supervision, P.C.

