## Supplemental Table for "Beach sand beneath our feet: an overlooked reservoir of antibiotic resistance genes and pathogens, revealed by metagenomic evidence from Qingdao’s recreational beaches": Supplementary table 1-2.docx

**Supplementary table 1.** General characteristics of samples, including number of valid tags, average length of valid tags and number of predicted OTUs.

| **Sample_ID** | **valid_tags** | **valid meanLength** | **OTU_counts** |
| --- | --- | --- | --- |
| SWZ1 | 61082 | 417.09 | 2529 |
| SWZ2 | 58899 | 417.25 | 2314 |
| SWZ3 | 60877 | 417.31 | 2393 |
| SWF1 | 65759 | 406.98 | 1623 |
| SWF2 | 64523 | 406.24 | 1615 |
| SWF3 | 64034 | 406.41 | 1759 |
| SWS1 | 66097 | 405.84 | 1627 |
| SWS2 | 66787 | 406.13 | 1726 |
| SWS3 | 65758 | 405.76 | 1583 |
| SWG1 | 64767 | 406.5 | 1657 |
| SWG2 | 65404 | 406.97 | 1871 |
| SWG3 | 66386 | 407.29 | 1860 |
| WWZ1 | 59612 | 420.65 | 994 |
| WWZ2 | 60164 | 421.73 | 945 |
| WWZ3 | 57784 | 418.1 | 1077 |
| WWF1 | 57561 | 420.05 | 2894 |
| WWF2 | 59951 | 420.52 | 3008 |
| WWF3 | 57914 | 420.16 | 2913 |
| WWS1 | 59193 | 423.99 | 1265 |
| WWS2 | 58391 | 424.21 | 1149 |
| WWS3 | 56820 | 424.03 | 1293 |
| WWG1 | 59017 | 419.27 | 1760 |
| WWG2 | 61092 | 419.03 | 1668 |
| WWG3 | 58027 | 419.14 | 1591 |
| SSZ1 | 54792 | 416.69 | 4261 |
| SSZ2 | 52106 | 416.87 | 4289 |
| SSZ3 | 53903 | 416.28 | 4246 |
| SSF1 | 56132 | 414.46 | 4255 |
| SSF2 | 57656 | 414.23 | 4512 |
| SSF3 | 60470 | 414.93 | 4459 |
| SSS1 | 57248 | 417.62 | 4882 |
| SSS2 | 58480 | 417.66 | 4893 |
| SSS3 | 59142 | 417.61 | 4823 |
| SSG1 | 63160 | 417.71 | 4668 |
| SSG2 | 57022 | 418.02 | 4743 |
| SSG3 | 58486 | 418.37 | 4801 |
| WSZ1 | 55657 | 414.21 | 2591 |
| WSZ2 | 54332 | 414.83 | 2683 |
| WSZ3 | 55020 | 414.25 | 2602 |
| WSF1 | 63007 | 413.27 | 3546 |
| WSF2 | 65274 | 413.44 | 4395 |
| WSF3 | 61704 | 413.83 | 3529 |
| WSS1 | 60569 | 420.96 | 2604 |
| WSS2 | 61618 | 421.33 | 2234 |
| WSS3 | 61666 | 421.83 | 2132 |
| WSG1 | 58519 | 417.36 | 4243 |
| WSG2 | 58297 | 417.07 | 4216 |
| WSG3 | 57352 | 417.31 | 4151 |

**Supplementary table 2.** Alpha diversity index statistical table.

| Samples | chao1 | observed_species | PD_whole_tree | shannon | simpson |
| --- | --- | --- | --- | --- | --- |
| SWZ | 3329.843 | 2311.832 | 103.85 | 6.24 | 0.9 |
| SSZ | 5482.236 | 4238.5 | 162.471 | 9.384 | 0.99 |
| WSZ | 3667.88 | 2606.4 | 110.721 | 7.9 | 0.98 |
| WWZ | 1494.969 | 966.967 | 48.823 | 5.307 | 0.92 |
| SWF | 2438.689 | 1538.101 | 72.122 | 4.948 | 0.821 |
| SSF | 5344.254 | 4318.532 | 163.328 | 9.82 | 1 |
| WSF | 4547.882 | 3679.1 | 146.786 | 9.958 | 1 |
| WWF | 3521.092 | 2874.899 | 126.764 | 8.86 | 0.99 |
| SWS | 2428.614 | 1498.6 | 71.619 | 4.659 | 0.79 |
| SSS | 5816.62 | 4766.866 | 176.759 | 10.162 | 1 |
| WSS | 3166.049 | 2214.134 | 95.855 | 7.553 | 0.98 |
| WWS | 1785.007 | 1201.467 | 55.9 | 5.826 | 0.92 |
| SWG | 2652.767 | 1649.866 | 78.32 | 5.296 | 0.86 |
| SSG | 5594.185 | 4627.601 | 172.458 | 10.177 | 1 |
| WSG | 5061.562 | 4118.833 | 159.934 | 9.577 | 0.99 |
| WWG | 2242.517 | 1621.467 | 74.435 | 7.34 | 0.98 |
